# Structural basis of RNA polymerase II transcription on the H3-H4 octasome

**DOI:** 10.1101/2025.05.13.653634

**Authors:** Cheng-Han Ho, Kayo Nozawa, Masahiro Nishimura, Mayuko Oi, Tomoya Kujirai, Mitsuo Ogasawara, Haruhiko Ehara, Shun-ichi Sekine, Yoshimasa Takizawa, Hitoshi Kurumizaka

## Abstract

The H3-H4 octasome is a nucleosome-like particle in which two DNA gyres are wrapped around each H3-H4 tetramer disk, forming a clamshell-like configuration. In the present study, we performed in vitro RNAPII transcription assays with the H3-H4 octasome and found that RNAPII transcribed the H3-H4 octasome more efficiently than the nucleosome. RNAPII paused at only one position, superhelical location (SHL) -4 in the H3-H4 octasome, in contrast to pausing at the SHL(−5), SHL(−2), and SHL(−1) positions in the nucleosome. Cryo-electron microscopy analysis revealed that two H3-H4 tetramer disks are retained when the RNAPII paused at the SHL(−4) position of the H3-H4 octasome. However, when RNAPII reached the SHL(−0.5) position, five base pairs before the dyad position of the H3-H4 octasome, the proximal H3-H4 tetramer was disassembled but the distal H3-H4 tetramer still remained on the DNA. Therefore, RNAPII efficiently transcribes the H3-H4 octasome by stepwise H3-H4 tetramer disassembly.

## INTRODUCTION

In eukaryotic cells, genomic DNA is packed into the cell nucleus by chromatin, primarily through the formation of a fundamental unit called the nucleosome (1). Each nucleosome consists of DNA wrapped around a histone octamer, a specialized structure composed of two copies each of the core histones H2A, H2B, H3, and H4 (2) (Figure 1A and C). The locations of the DNA wrapped around the nucleosome are referred to as superhelical locations (SHLs) (2, 3) (Figure 1A). At the dyad axis of the nucleosome, the minor groove of the DNA facing outwards from the histone core is defined as SHL(0). Subsequent outward-facing minor grooves are referred to as SHL ±1-7, with the plus and minus signs indicating the direction. The distance between each SHL roughly corresponds to 10 base pairs of DNA.

**Figure 1.**
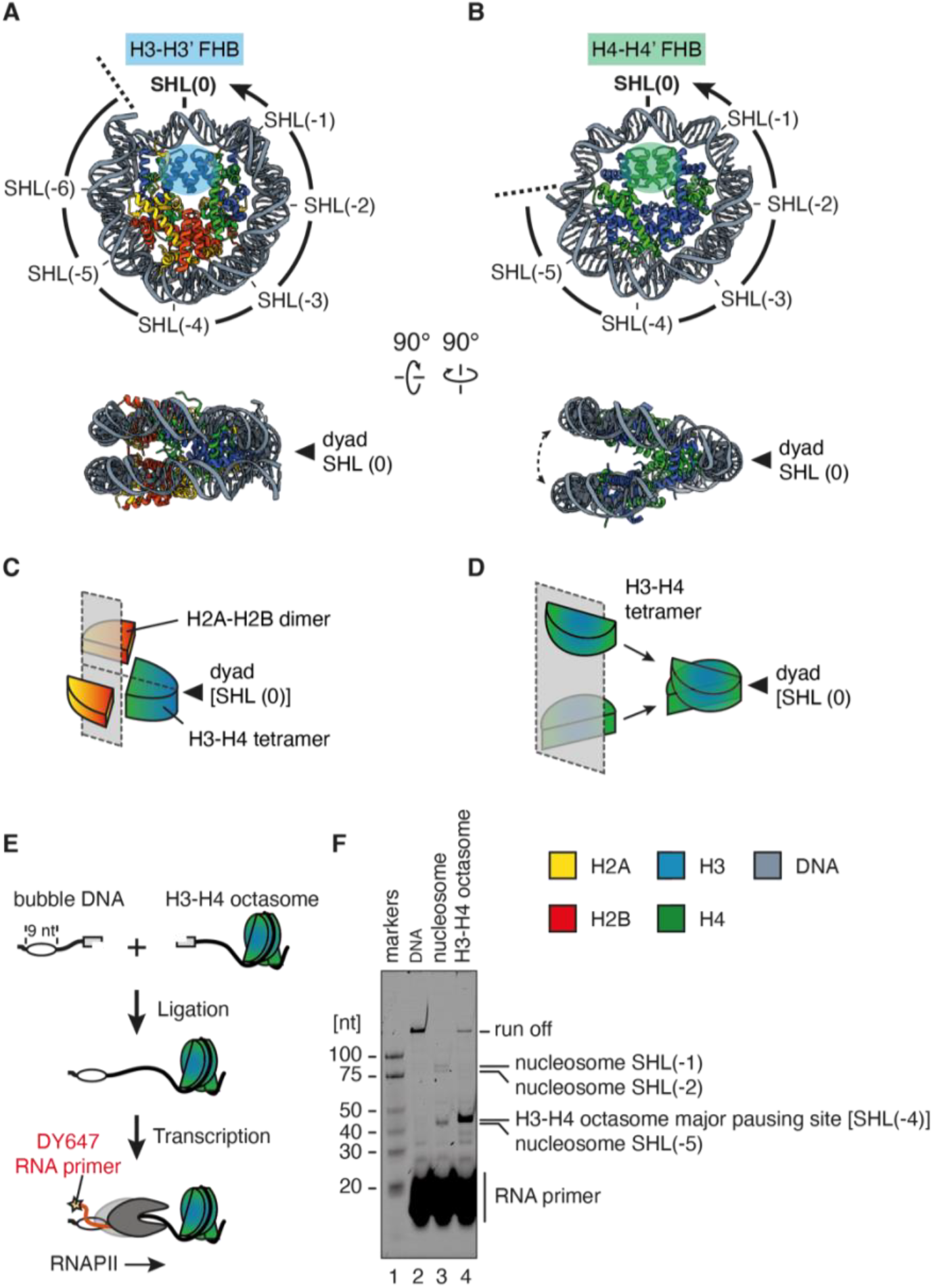
Transcription profile of the H3-H4 octasome. (**A**) Ribbon model of the nucleosome (PDB: 3LZ0), with the superhelical locations (SHLs) and the H3-H3′four-helix bundle (FHB) depicted. (**B**) Ribbon model of the H3-H4 octasome (closed form, PDB: 7×57), with the SHLs and the H4-H4′FHB depicted. (**C**) Schematic representation showing the relative positions of the H2A-H2B dimers and the (H3-H4)_2_ tetramer in the nucleosome. (**D**) Schematic representation showing the relative positions of the (H3-H4)_2_ tetramers in the H3-H4 octasome. (**E**) Schematic representation of the transcription assay. (**F**) Transcription assay with the nucleosome and the H3-H4 octasome. Denaturing gel electrophoresis analysis of the RNA products from the transcription assay with RNAPII. The assay was repeated three times independently and consistent results were obtained (Supplementary Figure S2).

Although the nucleosome contributes to packaging DNA in the nucleus, it also serves as functional machinery for many biological processes, such as replication (4), recombination (5), repair (6, 7), and transcription (8–18), as revealed by cryo-electron microscopy (cryo-EM). For transcription elongation, the DNA wrapped around the nucleosome is gradually peeled off by RNA polymerase II (RNAPII), while pausing at several superhelical locations: SHL(−6), SHL(−5), SHL(−2), and SHL(−1) (8, 10) (Figure 1A). Each of those SHLs are major DNA-histone interaction sites, suggesting that the DNA-histone interaction hinders the process of DNA-peeling by RNAPII. During the transcription of the nucleosome, the histone core remains octameric until RNAPII reaches SHL(−1). The nucleosome is then disassembled after the RNAPII passes through the SHL(0) position, and reassembled behind the RNAPII (15).

The nucleosome is a highly dynamic structure and may undergo transitions through various intermediate states, also known as subnucleosomal forms (19–21). Subnucleosomes are histone-DNA complexes with different stoichiometries compared to the nucleosome. The H3-H4 octasome is a subnucleosome, in which the DNA is wrapped around the H3-H4 octamer with two (H3-H4)_2_ tetramers (22–25) (Figure 1B and D). Early electron microscopy studies and a previous atomic force microscopy analysis demonstrated that the H3-H4 octasome forms a bead-like structure with dimensions similar to that of the nucleosome (26, 27). Our recent cryo-EM analysis further revealed the detailed structures of the H3-H4 octasome (25). As in the nucleosome, the DNA is left-handedly wrapped around the H3-H4 histone octamer, but significant differences exist between the H3-H4 octasome and the nucleosome. First, the histone positioning relative to SHL(0) is different. In the nucleosome, the H3-H3′ four helix bundle (FHB) (3), the interaction site of the two H3 protomers in the (H3-H4)_2_ tetramer, is located at SHL(0) (Figure 1A). In contrast, in the H3-H4 octasome, the H3-H3′FHBs are located at the SHL(±3) positions, flanking the SHL(0) (dyad), and a novel H4-H4′FHB is present at SHL(0) (Figure 1B). Second, approximately 120 base pairs of DNA are wrapped around the H3-H4 octasome, which is shorter than the 145-147 base pairs of DNA wrapped in the nucleosome (2) (Figure 1A and B). Third, two (H3-H4)_2_ tetramers of the H3-H4 octasome form a clamshell-like conformation, which is flexible and expands the distance between two DNA gyres compared to the nucleosome (Figure 1B and D).

The unique features of the H3-H4 octasome, particularly in terms of histone positioning, DNA wrapping length, and clamshell-like structural flexibility, suggest potential implications for distinct transcriptional regulation mechanisms. To study the transcription profile of the H3-H4 octasome by RNAPII, in the present study, we performed *in vitro* nucleosome transcription experiments coupled with cryo-EM analysis.

## MATERIAL AND METHODS

### Purification of proteins

Human histones H3.1 and H4 were prepared as previously described (28). Briefly, N-terminally hexa-histidine (His_6_)-tagged H3.1 was expressed in *E. coli* BL21 (DE3) cells, and N-terminally His_6_-tagged H4 was expressed in *E. coli* JM109 (DE3) cells. The *E. coli* cells were then sonicated and the lysates were denatured. Under denaturing conditions, the proteins were purified by nickel-nitrilotriacetic acid agarose (Ni-NTA) resin (QIAGEN) chromatography. The His_6_-tag was removed by thrombin protease under non-denaturing conditions. Finally, the histones were purified by cation exchange chromatography under denaturing conditions on a MonoS column. The collected histone fractions were dialyzed against water and lyophilized. RNAPII was purified from the yeast *Komagataella pastoris*, and TFIIS was purified as a recombinant protein as previously described (29).

### Preparation of DNA fragment for H3-H4 octasome reconstitution

Two kinds of 153 base pair DNA fragments were prepared for reconstituting H3-H4 octasomes.

To reconstitute the H3-H4 octasome initially used in the transcription assays, the DNA fragment (unmodified) was prepared as previously described (8). Briefly, the plasmid DNA containing a modified version of the Widom 601 sequence (30) was amplified in *E. coli*. The plasmid DNA was then collected and digested by *Eco*RV to obtain the target DNA fragment. After purification by polyethylene glycol precipitation, the DNA fragment was dephosphorylated to prevent self-ligation in the following steps. Finally, the DNA fragment was cleaved by *Bgl*I (Takara) and purified by DEAE-5PW anion-exchange column chromatography (TOSOH). The sequence is identical to the previously used DNA fragment (8).

To reconstitute the modified H3-H4 octasome used in the subsequent transcription assays and cryo-EM analysis, the DNA fragment (modified) was prepared by PCR. The modified DNA fragment is similar to the unmodified version, with only ‘ATT’ substituted with ‘TAA’ in the template strand (the resulting sequence is shown at the end of the paragraph). After PCR, the DNA fragment was then dephosphorylated and cleaved by *Bgl*I. Finally, using a Prep Cell apparatus (Bio-Rad), the DNA was purified by non-denaturing polyacrylamide gel electrophoresis. The resulting sequence is as follows, with the modified locations shown in bold letters:

Template strand: 5′-ATCAG AATCC CGGTG CCGAG GCCGC TCAAT TGGTC GTAGA CAGCT CTAGC ACCGC TTAAA CGCAC GTACG CGCTG TCCCC CGCGT TTTAA CCGCC AAGGG G**TAA**A CACCC AAGAC ACCAG GCACG AGACA GAAAA AAACA ACGAA AACGG CCACC A-3′;

Non-template strand: 5′-TGGCC GTTTT CGTTG TTTTT TTCTG TCTCG TGCCT GGTGT CTTGG GTGT**T TA**CCC CTTGG CGGTT AAAAC GCGGG GGACA GCGCG TACGT GCGTT TAAGC GGTGC TAGAG CTGTC TACGA CCAAT TGAGC GGCCT CGGCA CCGGG ATTCT GAT -3′.

### Preparation of the template H3-H4 octasomes containing 9 base-mismatched DNA region

The template H3-H4 octasome for transcription was prepared in a 2-step method, similar to the previously reported method for preparing the nucleosome template (8).

First, using the salt dialysis method, H3-H4 octasomes were reconstituted with one of the 153 bp DNA fragments described in the previous section (unmodified or modified) and the H3-H4 tetramer, which was refolded and purified as described previously (25). The H3-H4 octasomes were heated and purified by non-denaturing polyacrylamide gel electrophoresis, using a Prep Cell apparatus (Bio-Rad) (25).

Second, the short DNA fragment with a 9-base mismatch was ligated to the H3-H4 octasome by T4 DNA ligase (NIPPON GENE). The DNA sequence is identical to the one used in the previous study (8). After ligation, the H3-H4 octasomes were purified one final time by non-denaturing polyacrylamide gel electrophoresis, using a Prep Cell apparatus (Bio-Rad).

### Transcription assay on the H3-H4 octasomes

*In vitro* transcription assays using the unmodified template were performed by adding 0.1 μM RNAPII, 0.1 μM TFIIS and 0.4 μM primer RNA (5′-DY647-AUAAUUAGCUC-3′) (Dharmacon) to the reaction solution (26 mM HEPES-KOH (pH 7.5), 5 mM MgCl_2_, 50 mM potassium acetate, 0.3 μM zinc acetate, 33 μM Tris(2-carboxyethyl)phosphine, 1.6% glycerol, 400 μM UTP, 400 μM CTP, 400 μM GTP, and 400 μM ATP). To start the transcription reaction, 0.15 μM (final concentration) of template DNA, template nucleosome or template H3-H4 octasome was added to the reaction solution, in a final volume of 15 μL. The solution was then incubated at 30°C for 5 minutes. Subsequently, 4 μL of the reaction solution was collected and mixed with 2 μL of ProK solution (20 mM Tris (pH 7.5),20 mM EDTA,446 μg/μl proteinase K) to terminate the reaction. To denature the RNA product, 24 μL of Hi-Di formamide (Applied Biosystems) was added and the solution was incubated at 95°C for 5 minutes. Finally, the RNA products were fractionated by 10% denaturing polyacrylamide gel electrophoresis and detected by DY647 florescence, using an Amersham Typhoon scanner (Cytiva). DynaMarker DIG-labeled Blue Color Marker for Small RNA (BioDynamics Laboratory) served as the standard RNA markers.

*In vitro* transcription assays using the modified template were performed by mixing 0.225 μM naked DNA or H3-H4 octasome, 0.4 μM RNAPII, 0.15 μM TFIIS, and 0.6 μM primer RNA (5′-DY647-AUAAUUAGCUC-3′) (Dharmacon) in 10 μL of reaction solution (24 mM HEPES-KOH (pH 7.5), 5 mM MgCl_2_, 30 mM potassium acetate, 0.2 μM zinc acetate, 20 μM Tris(2-carboxyethyl)phosphine, 1% glycerol, 400 μM UTP, 400 μM CTP, 400 μM GTP, and 400 μM ATP or 40 μM 3 - ’ dATP). The solution was then incubated at 30°C. At the indicated times, 1 μL of the reaction solution was removed and mixed with 1 μL of ProK solution (20 mM Tris (pH 7.5), 20 mM EDTA, 446 μg/μL proteinase K) to terminate the reaction. To denature the RNA product, 24 μL of Hi-Di formamide (Applied Biosystems) was added and the solution was incubated at 95°C for 10 minutes. Finally, the RNA products were fractionated by 10% denaturing polyacrylamide gel electrophoresis and detected by DY647 florescence, using an Amersham Typhoon scanner (Cytiva). DynaMarker DIG-labeled Blue Color Marker for Small RNA (BioDynamics Laboratory) served as the standard RNA markers.

### Preparation of the RNAPII-H3-H4 octasome/tetrasome complexes for cryo-EM analysis

The transcription reaction was conducted by mixing 0.23 μM H3-H4 octasome, 0.40 μM RNAPII, 0.15 μM TFIIIS, and 0.6 μM primer RNA (5′-DY647-AUAAUUAGCUC-3′) (Dharmacon) in 1,000 mL of reaction solution (33 mM HEPES-KOH (pH 7.5), 3 mM Tris-HCl (pH 7.5), 5.0 mM MgCl_2_, 400 μM UTP, 400 μM CTP, 400 μM GTP, 400 μM ATP, 97 mM KoAC, 0.64 μM Zn(OAc)_2_, 64 μM TCEP-HCl, 0.15 mM DTT, and 3.2% glycerol). The solution was incubated at 30°C for 40 minutes, and then 20 μL of 500 mM EDTA was added to terminate the reaction. The complex was further purified by the GraFix method (31), with low buffer composed of 20 mM HEPES-KOH (pH 7.5), 50 mM KOAc, 0.2 μM Zn(OAc)_2,_ 0.5 mM TCEP-HCl, and 10% (w/v) sucrose, and high buffer composed of 20 mM HEPES-KOH (pH 7.5), 50 mM KOAc, 0.2 μM Zn(OAc)_2,_ 0.5 mM TCEP-HCl, 25% (w/v) sucrose, and 0.1% glutaraldehyde. The reaction mixture was loaded onto the gradient, and the samples were centrifuged at 30,000 rpm and 4°C for 14 hours, using a Beckman SW32 Ti rotor.

After centrifugation, 1 mL fractions were collected from the top of the gradient solution, using a pipette. To determine the range for collection, the fractions were analyzed by 4% non-denaturing PAGE with RNAPII-H3-H4 octasome/tetrasome complex detection by SYBR Gold, and 10% denaturing PAGE with elongated RNA product visualization by DY647 fluorescence, using an Amersham Typhoon imager (GE Healthcare). Fractions containing the elongated RNAPII-H3-H4 octasome/tetrasome complexes were collected, and then buffer exchanged on a PD10 column into 20 mM HEPES-NaOH (pH 7.5), 0.2 μM Zn(OAc)_2,_ and 0.1 mM TCEP-HCl. Finally, the sample was concentrated by an Amicon Ultra 100K filter to a final DNA concentration of 0.51 mg/mL.

For cryo-EM analysis, Quantifoil grids (R1.2/1.3 Cu 200 mesh, Quantifoil Micro Tools) were glow discharged for 1 min with a PIB-10 ION Bombarder (Vacuum Device Inc.), immediately before use. Subsequent processes were performed using a Vitrobot Mark IV (Thermo Fisher Scientific). Portions (2.5 μL) of diluted samples (DNA concentration 0.05 mg/mL) were applied to the grids, blotted for 4 or 5 seconds with a blot force of 5 at 4°C/100% humidity, and plunge-frozen.

### Cryo-EM data collection

Cryo-EM images were recorded by a Krios G4 transmission electron microscope (Thermo Fisher Scientific), equipped with a K3 direct electron detector (Gatan). Automated data acquisition was performed with the EPU software (pixel size: 1.06 Å, defocus values: between -1.0 and -2.5 μm, total dose: 59.2∼60.8 e^-^/A^2^ for over 40 frames). In total, three data sets were collected. More information can be found in *SI Appendix* Table 1.

### Cryo-EM image processing

The cryo-EM images of each data set were imported into Relion4 for single particle analysis (32). Motion correction was performed by MotionCor2 (33), and CTF estimation was performed by CTFFIND4 (34). Subsequently, micrographs with poor quality were discarded by manual selection.

A subset of 1,000 micrographs from the first data set were then chosen for training a Topaz model for auto-picking. In brief, auto-picking was first performed in the Laplacian-of-Gaussian mode, and 2D classification was performed to discard junk particles. The remaining particles were used for training the first Topaz model.

This Topaz model was then used for auto-picking particles from the entire first data set (35). 2D and 3D classifications were performed to discard junk particles. The remaining particles were then used to train a second Topaz model, which was the main model used for auto-picking particles from the first, second, and third data sets.

The particles from these different data sets were then merged and subjected to 2D and 3D classifications (Supplementary Figure S4). In the 2^nd^ round of 3D classification, two major classes of the particles can be observed. These two classes were selected separately, and all downstream analyses of the two classes were performed independently.

For the SHL(−4)-like particles, several additional rounds of 3D classification were performed in Relion to remove bad particles and eliminate the effect of orientation bias (Supplementary Figure S5). The remaining good particles were imported to CryoSPARC (36) for subsequent processing. To remove particles in which the H3-H4 octasome subunit was unclear, one extra round of 3D classification (without alignment) was performed in CryoSPARC with an H3-H4 octasome mask. Finally, Bayesian-polished particles of the chosen class were subjected to 3D flexible refinement. The output map of the 3D flexible refinement was sharpened for model building and is displayed in Figure 2.

**Figure 2.**
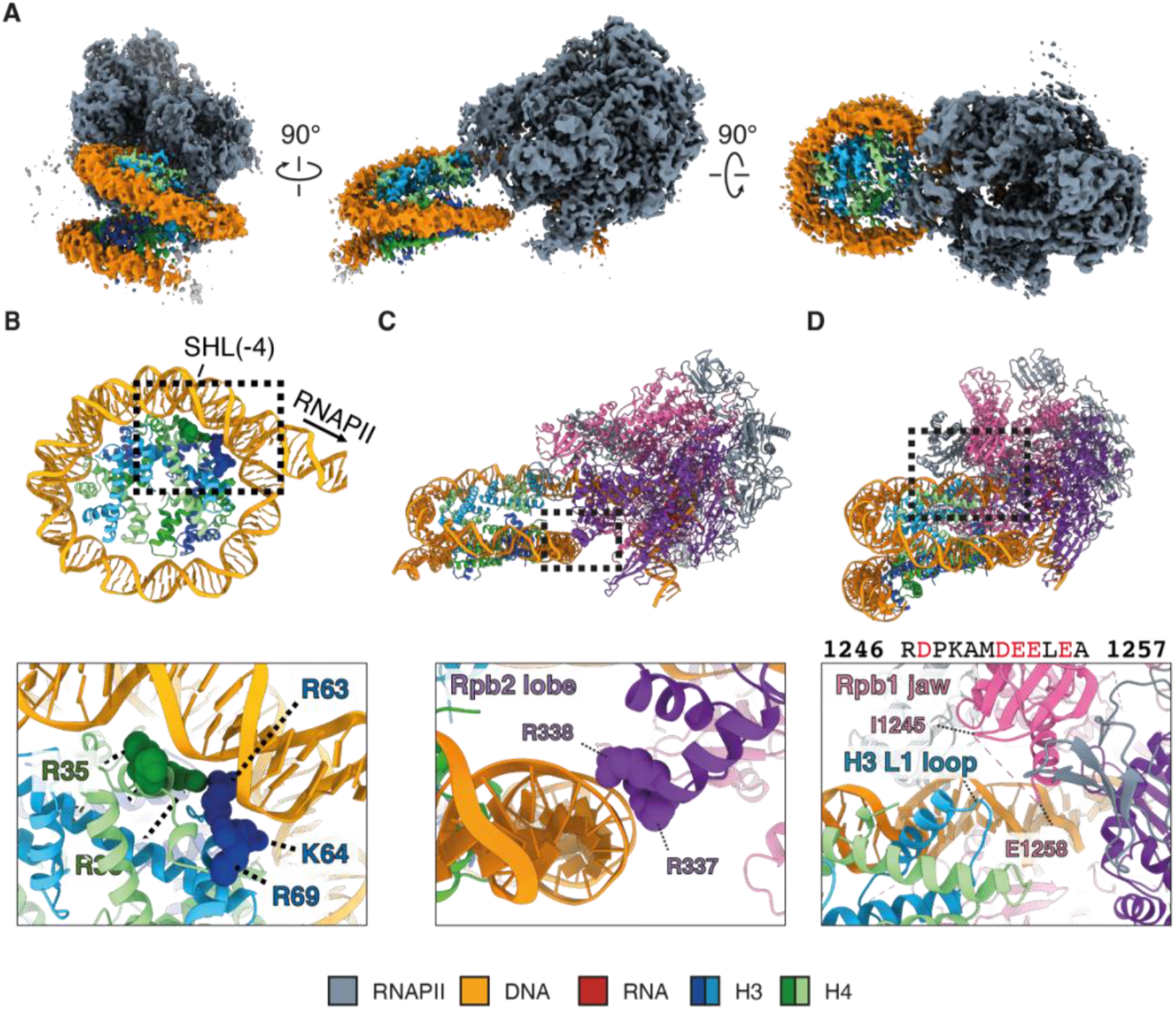
Cryo-EM structure of the RNAPII-H3-H4 octasome [SHL(−4)]. (**A**) Cryo-EM density maps of the RNAPII-H3-H4 octasome [SHL(−4)]. Three different views of the same map are shown. (**B**) Interactions between DNA and histones when RNAPII pauses at SHL(−4) in the H3-H4 octasome. The lower panel shows a close-up view of the boxed region in the upper panel. The histone residues of H3 (blue) and H4 (green) that interact with DNA are indicated. (**C**) Interaction between RNAPII subunit Rpb2 (purple) and H3-H4 octasomal DNA (orange). The lower panel shows a close-up view of the boxed region in the upper panel. The arginine residues of Rpb2 involved in the interaction are indicated. (**D**) Possible interaction between RNAPII subunit Rpb1 (pink) and the H3 L1 loop (blue). The lower panel shows a close-up view of the boxed region in the upper panel. The amino acid residues between I1245 and E1258 of the Rbp1 jaw, which are invisible in this structure, are shown above the lower panel. Several basic residues in this region could interact with the H3 L1 loop, which contains several acidic residues.

For the SHL(−0.5)-like particles, one additional round of 3D classification was performed in Relion, before importing the selected particles into CryoSPARC (Supplementary Figure S6). At this stage, we noticed that severe orientation bias was present in the SHL(−0.5) particles, making it impossible to obtain a clear map of both the RNAPII and the H3-H4 tetrasome. Because the structure of RNAPII is not expected to differ much from previous reported structures, we decided to focus mainly on the H3-H4 tetrasome instead, which is the highlight of this study. To visualize the histone helices of the H3-H4 tetrasome, two rounds of 3D classifications (without alignment) were performed in CryoSPARC with an H3-H4 tetrasome mask. A subsequent 3D classification (without alignment) was performed to slightly reduce the effect of orientation bias on the RNAPII. It should be noted that the numerous rounds of 3D classification (in attempts to reduce the effect of orientation bias) forced us to reject a large number of particles. Finally, Bayesian-polished particles of the selected class were subjected to 3D flexible refinement. The output map of the 3D flexible refinement was sharpened for model building and is displayed in Figure 3. More information can be found in Supplementary Table 1.

**Figure 3.**
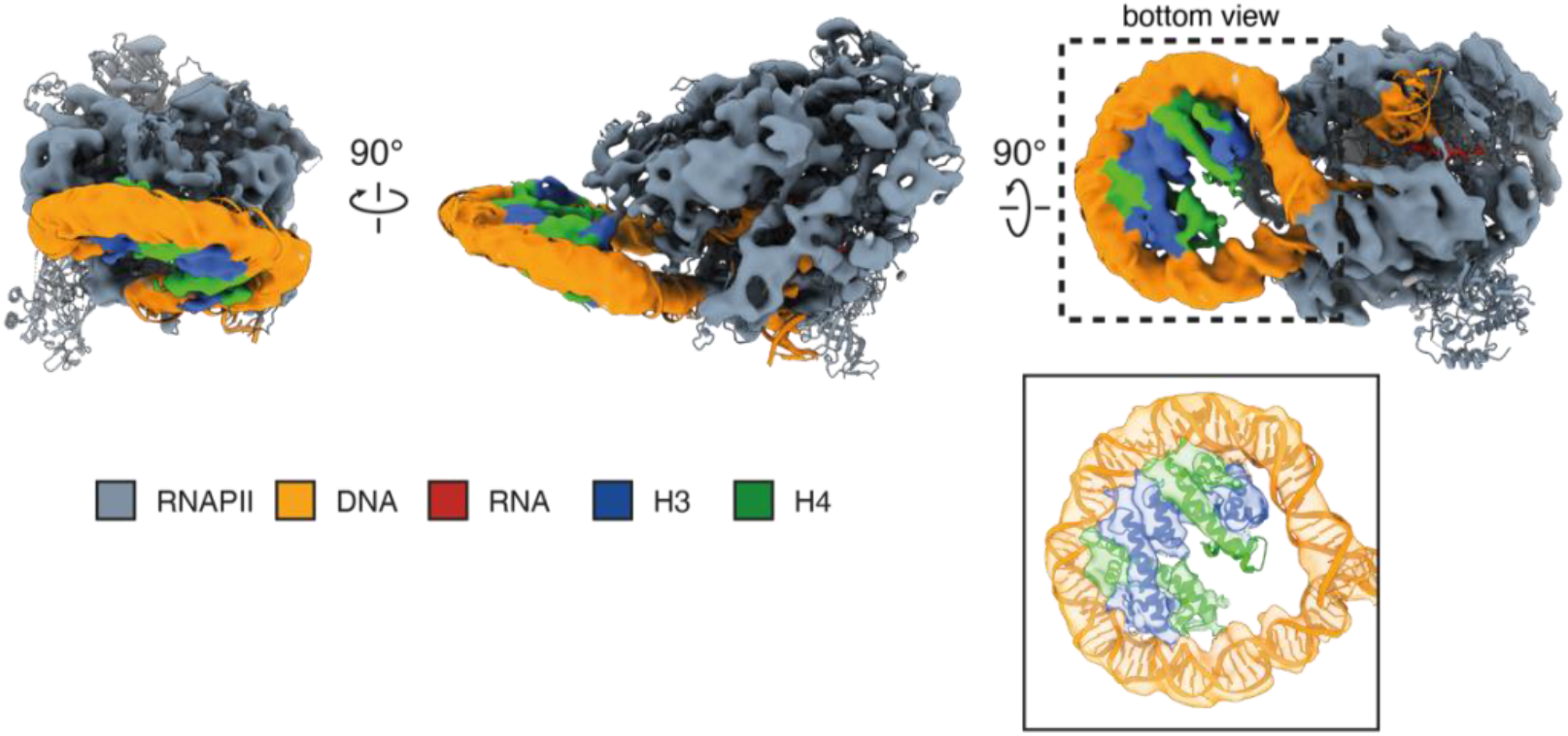
Cryo-EM structure of the RNAPII-H3-H4 tetrasome [SHL(−0.5)]. Three different views of the same cryo-EM density map (with the structure model fitted) are shown. One H3-H4 tetramer is retained in the structure, resulting in a H3-H4 tetrasome. Close-up view of the H3-H4 tetrasome region encircled with the box in the upper panel. The a-helices of H3 and H4 are visualized in the H3-H4 tetrasome.

### Model building

Modeling for both the RNAPII-H3-H4 octasome and the RNAPII-H3-H4 tetrasome complex was based on the atomic model of the RNAPII-nucleosome complex stalled at SHL(−5) (PDB: 6A5P) (8). For the RNAPII-H3-H4 octasome complex, the nucleosome was first replaced by the H3-H4 octasome (closed form) atomic model (PDB: 7×57) (25). The edited atomic model was then refined using the phenix.real_space_refine tool in the Phenix package (37), and modeled manually to fix outliers by using the Coot software (38, 39). For the RNAPII-H3-H4 tetrasome complex, the nucleosome was replaced by half of the H3-H4 octasome (closed form) atomic model (PDB: 7×57) (25). Due to the limited resolution of the cryo-EM map, no further refinement was performed (only validation). More information can be found in Supplementary Table 1.

### Data availability

The data that support this study are available from the corresponding authors upon reasonable request. The cryo-EM reconstructions and atomic models of the RNAPII-H3-H4 octasome complex and the RNAPII-H3-H4 tetrasome complex generated in this study have been deposited in the Electron Microscopy Data Bank and the Protein Data Bank, under the accession codes EMD-XXXXX and PDB ID XXXX, and EMD-XXXXX and PDB ID XXXX, respectively. The structures used in this study can be found in the Protein Data Bank under the accession codes: 6A5P and 7×57.

## RESULTS

### The transcription profile of the H3-H4 octasome is remarkably different from those of the nucleosome

To investigate the transcription process of the H3-H4 octasome, we first prepared it with a previously reported template DNA sequence for the transcription assay (8) (Supplementary Figures S1A and B). The template DNA, which contains a 9-base mismatch, allows the RNA primer to bind (Figure 1E). This RNA primer is extended by transcription elongation with RNAPII and thus serves as an indicator of the RNAPII pausing position on the transcribed template. We purified RNAPII from the yeast *Komagataella phaffii* and prepared the elongation factor TFIIS as a recombinant protein, as previously described (29). Using the prepared H3-H4 octasome template, RNAPII, and TFIIS, we performed a transcription assay *in vitro*. The reaction mixtures were analysed by denaturing polyacrylamide gel electrophoresis, and the transcribed RNAs were detected. In this assay, the template DNA and the template nucleosome were also included for comparison.

In the transcription assay with the naked DNA template, RNAPII pausing was barely observed, and most of the RNAs were run-off transcription products (Figure 1F, lane 2).

Consistent with the previous studies, with the nucleosome template, RNAPII pauses at SHL(−5), SHL(−2), and SHL(−1) were observed, and the run-off transcript was barely detected (Figure 1F, lane 3) (8). Interestingly, with the H3-H4 octasome template, an intense but somewhat slower migrating band was observed near the nucleosomal SHL(−5) pausing site (Figure 1F, lane 4). We hypothesized that the H3-H4 octasome may be positioned similarly to the nucleosome in the template DNA. According to the hypothesis, the major pausing site of the H3-H4 octasome would correspond to a position after SHL(−5), possibly SHL(−4).The run-off transcript was detected with the H3-H4 octasome but not the nucleosome, suggesting that RNAPII can transcribe the H3-H4 octasome more efficiently than the nucleosome (Figure 1F, lane 4). No other noticeable bands were detected between the band just above the SHL(−5) band and the run-off band. This suggests that, unlike nucleosomal transcription, which has several pausing sites, RNAPII pauses only at one major site located near the H3-H4 octasome entry site, which may correspond to SHL(−4).

### Cryo-EM structure of RNAPII paused on the SHL(−4) position of the H3-H4 octasome

To clarify the mechanism of RNAPII pausing on the H3-H4 octasome, we performed cryo-EM analysis. The template DNA used in this cryo-EM analysis was modified so that RNAPII would pause at SHL(−0.5) (5 base-pairs before the SHL(0) position) after passing the major SHL(−4) pausing site (Supplementary Figure S3A). Under the experimental setup, RNAPII pauses at the SHL(−0.5) site after overcoming the major natural pausing site, which may provide another snapshot structure during the transcription process by cryo-EM with one sample preparation (Supplementary Figures S3B and C). This allows us to trace the structural transition during the RNAPII passage through the H3-H4 octasome. Using this experimental setup, we purified the RNAPII-H3-H4 octasome complex by sucrose/glutaraldehyde gradient ultracentrifugation (GraFix) method (Supplementary Figure S3D) and proceeded to the cryo-EM single particle analysis.

We first obtained the high resolution cryo-EM structure of RNAPII paused at the major natural pausing site of the H3-H4 octasome, which indeed corresponds to the SHL(−4) position (Figure 2A and Supplementary Figures S4, S5, and S7A-C). RNAPII collides with the H3-H4 octasome at the major natural SHL(−4) pausing site, without affecting the histone containing two (H3-H4)_2_ tetramers. In nucleosome transcription, RNAPII pauses at the major contact sites between DNA and core histones(8). Consistently, in the H3-H4 octasome, RNAPII pauses on the DNA contacting 6 basic amino acid residues (H3R63, H3K64, H3R69, H4K31, H4R35, and H4R36) around the SHL(−4) position (Figure 2B). Interestingly, these residues also induce RNAPII pausing at the SHL(−1) position in the nucleosome (8).

In the RNAPII-H3-H4 octasome complex paused at the SHL(−4) position, two arginine residues R337 and R338 in the lobe of the RNAPII Rpb2 subunit are near the DNA wrapped around the H3-H4 octamer (Figure 2C). In addition, the RNAPII subunit, Rpb1, may contact the H3 L1 loop (Figure 2D). The R1246-A1257 region of the Rpb1 jaw contains several acidic residues, which may interact with the H3 L1 loop via the basic residues K79 and R83, although the Rpb1 R1246-A1257 region is not visible, probably due to its flexibility or multiple conformations (Figure 2D, lower panel). These histone-DNA contacts at the SHL(−4) position, the RNAPII-DNA interaction, and the RNAPII-histone interaction may promote the RNAPII pausing at the SHL(−4) position of the H3-H4 octasome.

### An (H3-H4)2 tetramer dissociates after RNAPII overcomes the SHL(−4) barrier

In the template DNA used for cryo-EM analysis, a ‘T’ site is newly added at the position 74 base pairs from the transcription priming site (presumably SHL(−0.5) position) of the H3-H4 octasome (Supplementary Figure S3A). A ‘TTTT’ site also exists 13 base pairs downstream of the inserted ‘T’ site. *In vitro* H3-H4 octasome transcription reactions were then conducted in the presence of 3 - ’ dATP, instead of ATP. A single ‘T’ insertion may be too weak to completely stop the RNAPII progression; therefore, the RNA product that originated from RNAPII stalled at the ‘TTTT’ site was also detected (Supplementary Figures S3B and C).

Using the cryo-EM sample prepared by the GraFix method from the previous section (Supplementary Figure S3D), we also determined the cryo-EM structure of RNAPII intentionally paused at the SHL(−0.5) position of the H3-H4 octasome, which corresponds to the inserted ‘T’ site. Note that we could not obtain the structure of RNAPII paused at the ‘TTTT’ site corresponding to the SHL(+1) position (Supplementary Figures S4, S6, and S7D-F). It is possible that when transcription has reached after passing the dyad of the H3-H4 octasome and paused at the ‘TTTT’ site, histones H3 and H4 have completely dissociated, leaving only the naked DNA.

In the SHL(−0.5) structure, RNAPII is paused five base pairs before the dyad SHL(0) position (Figure 3). The resolution of this structure is limited as compared to the SHL(−4) structure, possibly due to the small number of particles used in the final reconstruction. Strikingly, we noticed that only one (H3-H4)_2_ tetramer is present in the SHL(−0.5) structure (Figure 3). This result suggests that an (H3-H4)_2_ tetramer may have dissociated from the H3-H4 octasome after RNAPII surmounted the SHL(−4) barrier but before it reached the SHL(−0.5) position. Because the remaining H3-H4 tetrasome may be relatively unstable, this could explain why RNAPII does not pause after overcoming the natural barrier at the SHL(−4) position. Therefore, the two H3-H4 tetramer units may be dissociated in a stepwise manner from the H3-H4 octasome during RNAPII passage.

## DISCUSSION

We previously reported that the H3-H4 octasome wraps the DNA differently from the nucleosome (25). Although the H3-H4 octasome has been suggested to exist in cells, its function has remained elusive. In this study, we found that the transcription profile of the H3-H4 octasome differs from that of the nucleosome (Figure 1F). During nucleosome transcription, RNAPII pauses at several superhelical locations, including SHL(−6), SHL(−5), SHL(−2), and SHL(−1). These nucleosome barriers of RNAPII transcription are drastically reduced by elongation factors, such as Spt4/5 and Elf1, and the nucleosome is eventually transferred to the upstream DNA region with the aid of other elongation factors and the histone chaperone FACT(10, 15). In contrast, RNAPII transcription of the H3-H4 octasome pauses only at SHL(−4). After overcoming the SHL(−4) barrier, RNAPII does not pause and finishes transcribing the remaining template (Fig 1F and Movie S1). Given that RNAPII alone pauses at multiple positions in the nucleosome, the H3-H4 octasome may lead to more efficient transcription elongation than the nucleosome (Figure 4).

**Figure 4.**
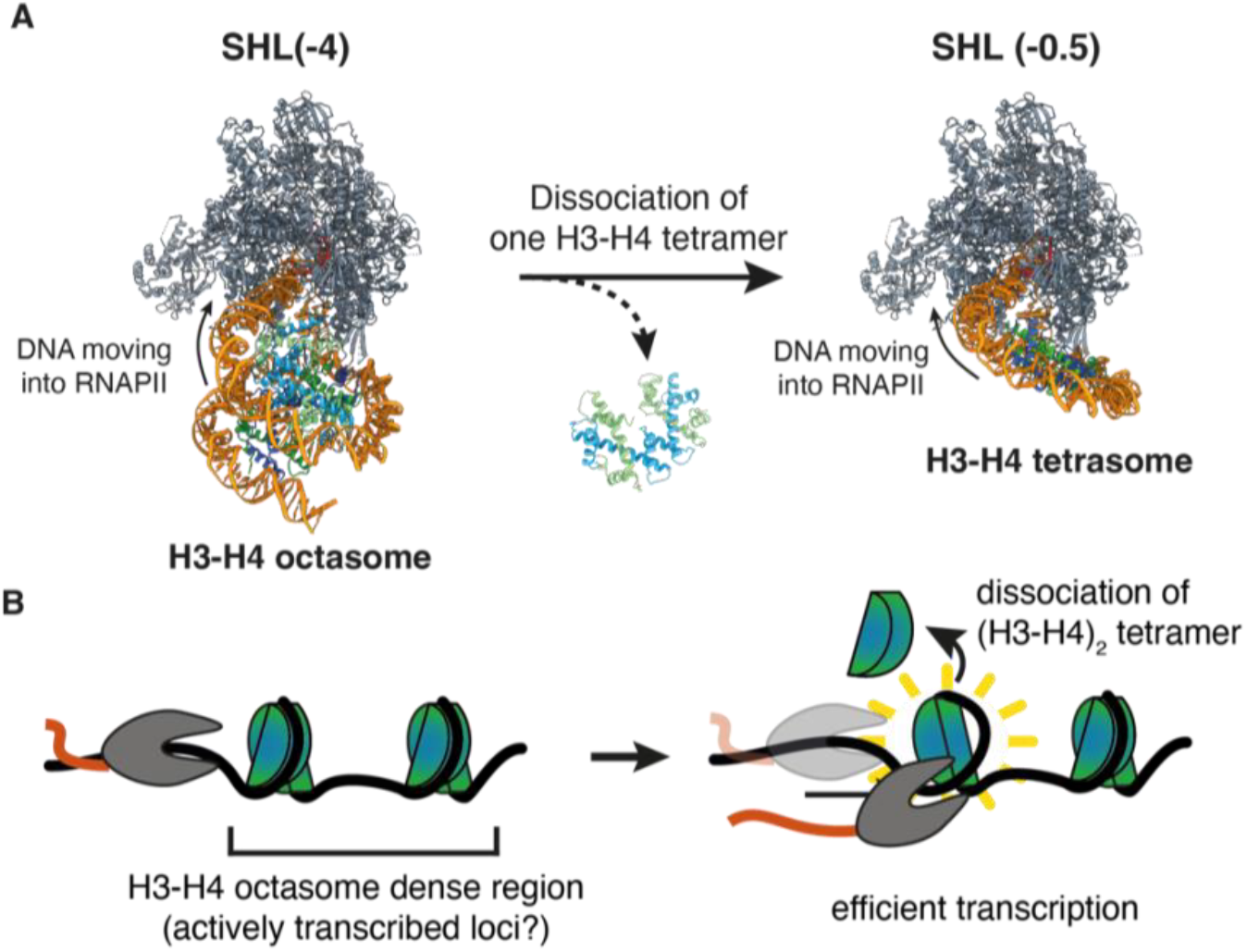
Model of transcription on the H3-H4 octasome. (*A*) Snapshots of the transcription process of the H3-H4 octasome. When RNAPII encounters the H3-H4 octasome SHL(−4) (left panel), it pauses due to the DNA-histone interactions within the H3-H4 octasome and the interactions between RNAPII subunits and the H3-H4 octasome. As RNAPII overcomes the SHL(−4) barrier, one H3-H4 tetramer dissociates from the H3-H4 octasome, leaving an H3-H4 tetrasome (right panel). This H3-H4 tetrasome is structurally unstable, and therefore incapable of stopping RNAPII. (*B)* In contrast to the nucleosomal transcription process, in which RNAPII pauses at SHL(−5), SHL(−2), and SHL(−1), in the H3-H4 octasomal transcription process, RNAPII only pauses at SHL(−4). This suggests that H3-H4 octasomes may be transcribed more efficiently than nucleosomes.

Nevertheless, conserved mechanisms of RNAPII pausing exist between the H3-H4 octasome and the nucleosome. The strong histone-DNA interactions within the H3-H4 octasome at the SHL(−4) position likely contribute to the natural RNAPII pausing (Figure 2). In the nucleosome, this histone-DNA interaction of the H3-H4 octasome is conserved in the SHL(−1) position, which also induces strong natural RNAPII pausing (8). In the nucleosome SHL(−1) position, the H3R63, H3K64, H3R69, and H4R36 residues interact with DNA, forming an obstacle for RNAPII to peel off the DNA. These histone residues also contribute to the SHL(−4) pausing in the H3-H4 octasome (Figure 2B), suggesting that the pausing mechanism induced by histone-DNA interactions may be conserved between the nucleosome and the H3-H4 octasome, and possibly also among other non-canonical nucleosomes. However, more investigation is required to determine whether the histone-DNA interaction is a universal mechanism in transcription pausing.

In summary, the present study reveals how a recently reported atypical nucleosome, the H3-H4 octasome, is transcribed by RNAPII. The fact that RNAPII only pauses at one site, SHL(−4), suggests that the H3-H4 octasome may be transcribed more efficiently than the nucleosome. Therefore, the H3-H4 octasome may exist in highly transcribed genomic loci.

Remaining issues to understand are as follows. (i) How do transcription elongation factors such as Spt4/5 and Elf1 affect the transcription of the H3-H4 octasome? (ii) Is the H3-H4 octasome reassembled or discarded after the RNAPII passes through its dyad DNA region? (iii) Where do the genomic loci enriched in H3-H4 predominantly exist and function? The H3-H4 octasome lacks the acidic patch, a region in the nucleosome where various chromatin-interacting factors, such as histone chaperones, nucleosome remodelers, and histone modifiers, typically bind. Therefore, the lack of acidic patch in the H3-H4 octasome may have distinct functions in genome regulation from the canonical nucleosome. Further studies will be required to solve these issues.

## DATA AVAILABILITY

The data that support this study are available from the corresponding authors upon reasonable request. The cryo-EM reconstructions and atomic models of the RNAPII-H3-H4 octasome complex and the RNAPII-H3-H4 tetrasome complex generated in this study have been deposited in the Electron Microscopy Data Bank and the Protein Data Bank, under the accession codes EMD-64225 and PDB ID 9UJS, and EMD-64226 and PDB ID 9UJT, respectively.

## SUPPLEMENTARY DATA

Supplementary Data are available at NAR online.

## AUTHOR CONTRIBUTIONS

Cheng-Han Ho: Investigation, Formal analysis, Visualization, Writing—original draft.

Kayo Nozawa: Conceptualization, Funding acquisition, Investigation.

Masahiro Nishimura: Investigation, Resources.

Mayuko Oi: Investigation, Resources.

Tomoya Kujirai: Formal analysis, Funding acquisition, Methodology.

Mitsuo Ogasawara: Investigation, Formal analysis.

Haruhiko Ehara: Resources. Shun-ichi Sekine: Resources.

Yoshimasa Takizawa: Formal analysis, Funding acquisition.

Hitoshi Kurumizaka: Conceptualization, Funding acquisition, Project administration, Supervision, Witing—review & editing.

## ACKNOWLEDGEMENTS

We would like to express our deepest gratitude to Y. Iikura, M. Dacher, and Y. Takeda (Univ. of Tokyo) for their assistance. ChatGPT was used for grammatical correction, rephrasing and/or rearranging parts of the text. No original content was produced by AI.

