## Supplementary Figures for "Structural basis of RNA polymerase II transcription on the H3-H4 octasome"

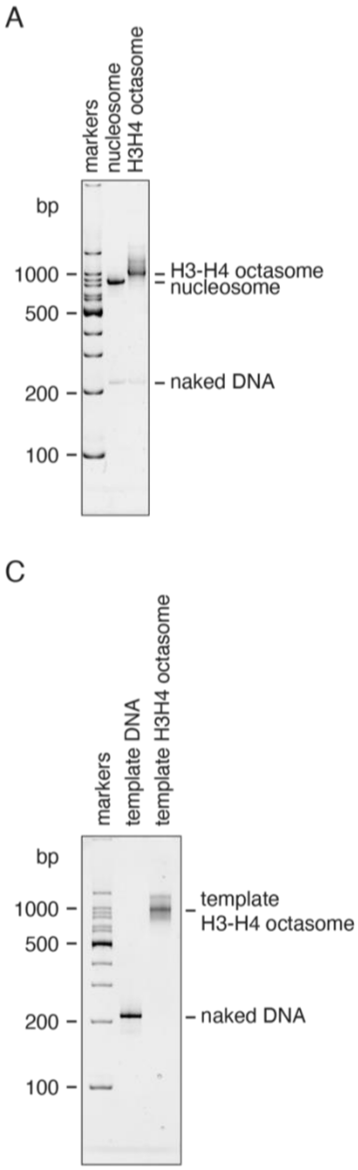


**Supplementary Figure S1. H3-H4 octasome sample preparations.** (A) Purified nucleosome and H3-H4 octasome (reconstituted with the unmodified DNA) were analyzed by native polyacrylamide gel electrophoresis (PAGE) and visualized by ethidium bromide (EtBr) staining to detect DNA. (B) The same samples from (A) were also analyzed by SDS-PAGE and visualized by Coomassie Brilliant Blue (CBB) staining to detect proteins. (C) Purified template DNA and H3-H4 octasome (reconstituted with the modified template DNA) were analyzed by native-PAGE and visualized by EtBr staining to detect DNA. (D) The H3-H4 octasome from (C) was also analyzed by SDS-PAGE and visualized by CBB staining to detect proteins.

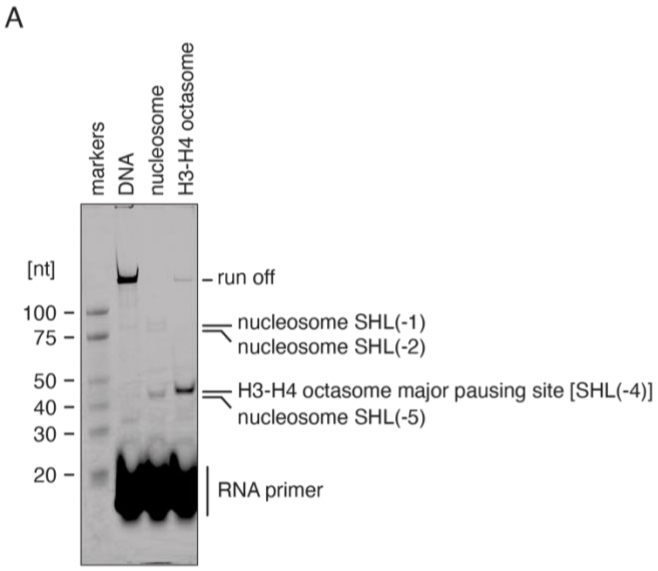


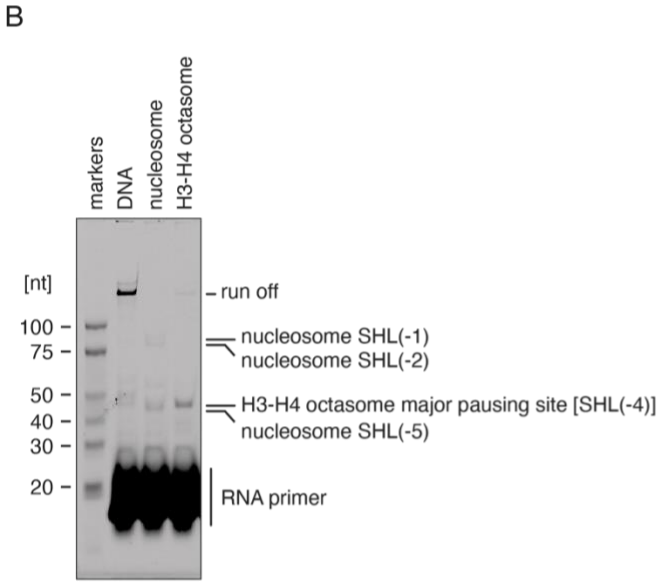


**Supplementary Figure S2. Transcription assay of the H3-H4 octasome using unmodified DNA template.** The transcription assay in Fig. 1f was repeated three times independently in total. The 2 repeats other than Fig. 1F is shown in (A) and (B).

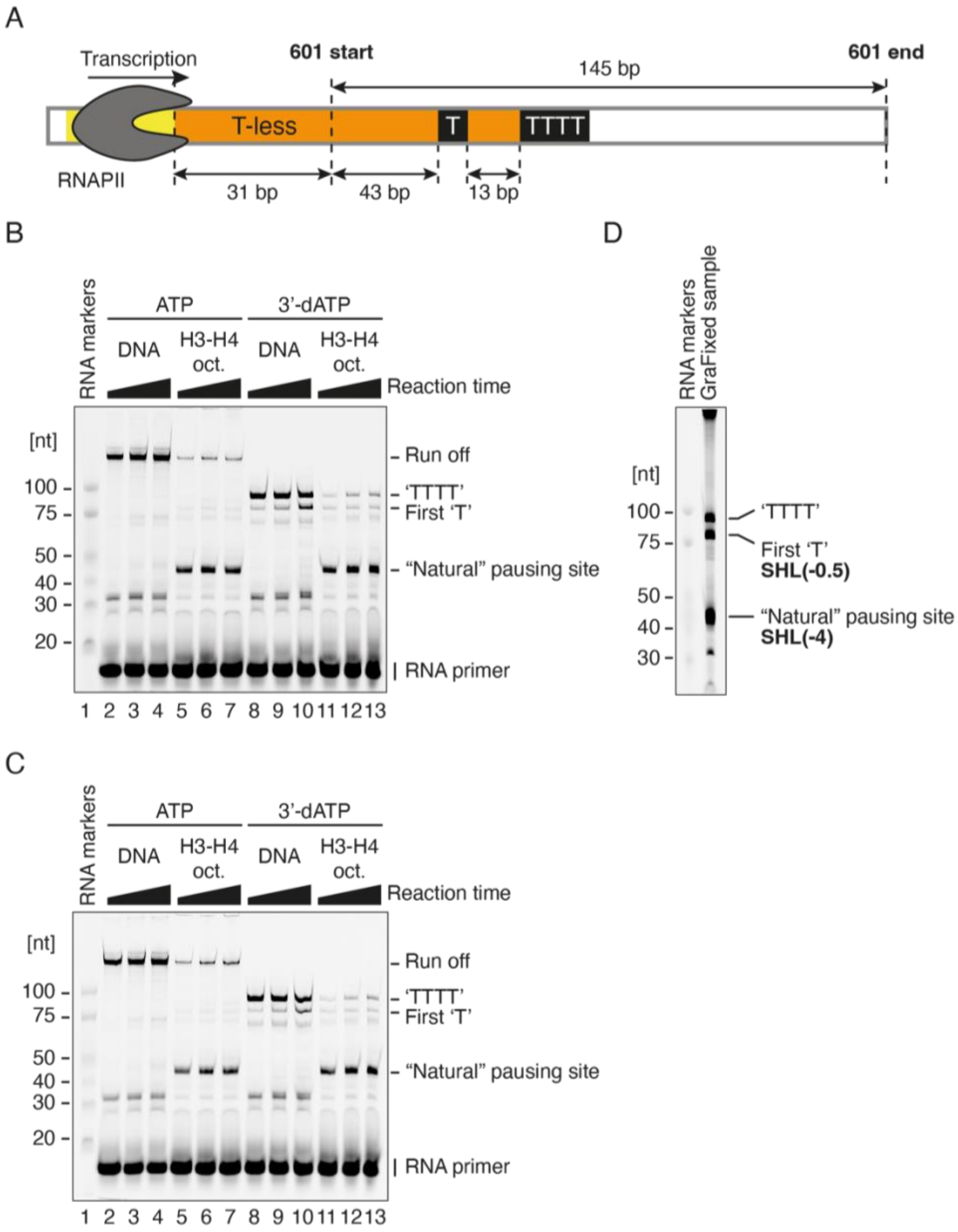


**Supplementary Figure S3. Transcription assay of the H3-H4 octasome using modified DNA template.** (A) The modified DNA template. A new 'T' site is added 74 bp downstream of the transcription start site. The original 'TTTT' site also exists 13 bp downstream of the inserted 'T' site. (B, C) Transcription assays using the modified DNA template. The reaction times were 5, 20, and 40 minutes. ATP or 3´-dATP was used as indicated. (D) The GraFix-processed RNAPII-H3-H4 octasome sample was analyzed by denaturing gel electrophoresis. The RNA products were detected.

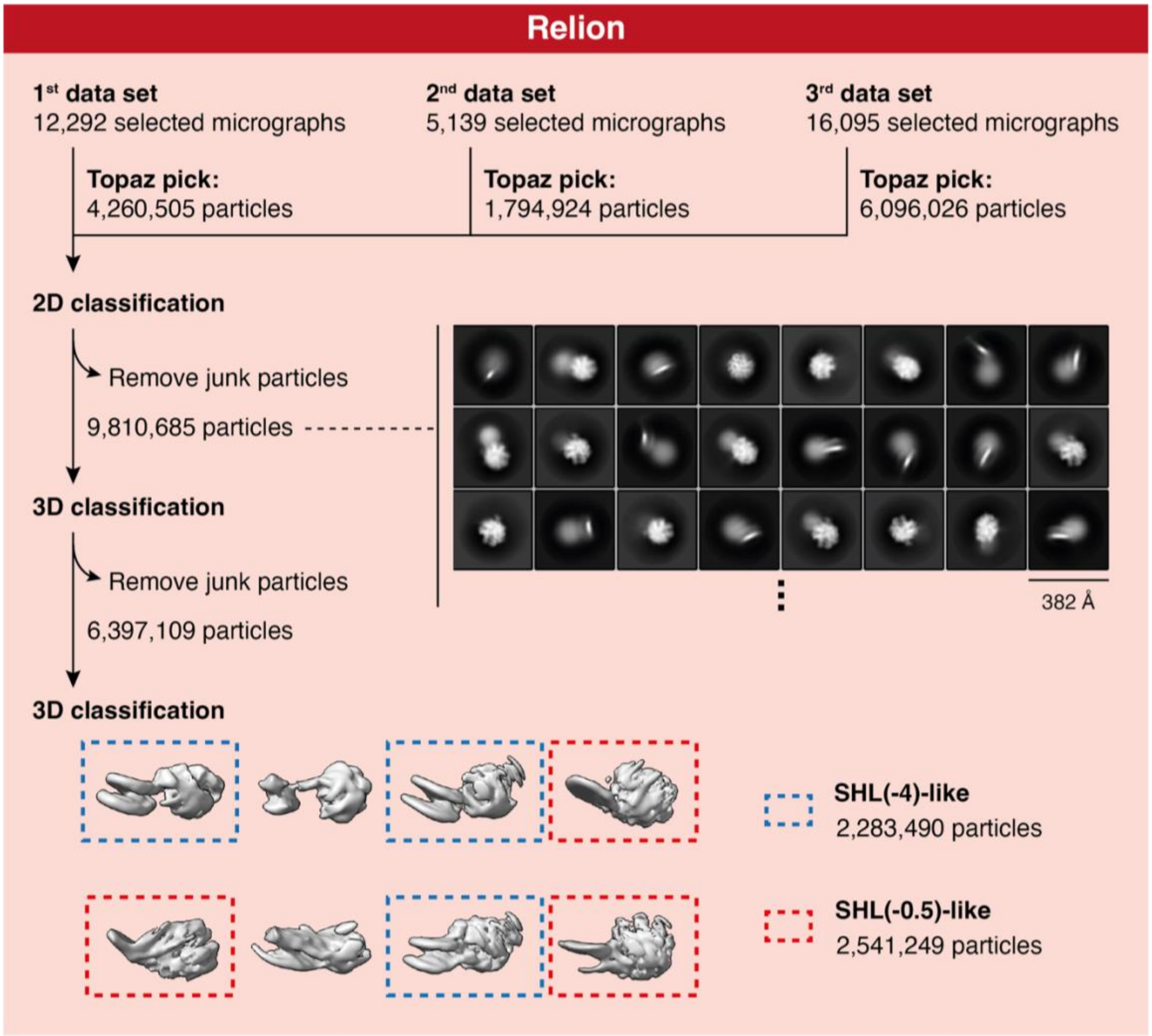


**Supplementary Figure S4. Cryo-EM single particle analysis of data sets by Relion.** Three data sets were obtained and analyzed independently before 2D classification. After Topaz picking, the particles were merged and subjected to 2D classification. A subset of the 2D classes is shown. After two rounds of 3D classification, SHL(-4)-like particles and SHL(-0.5)-like particles were separated and subsequently processed individually.

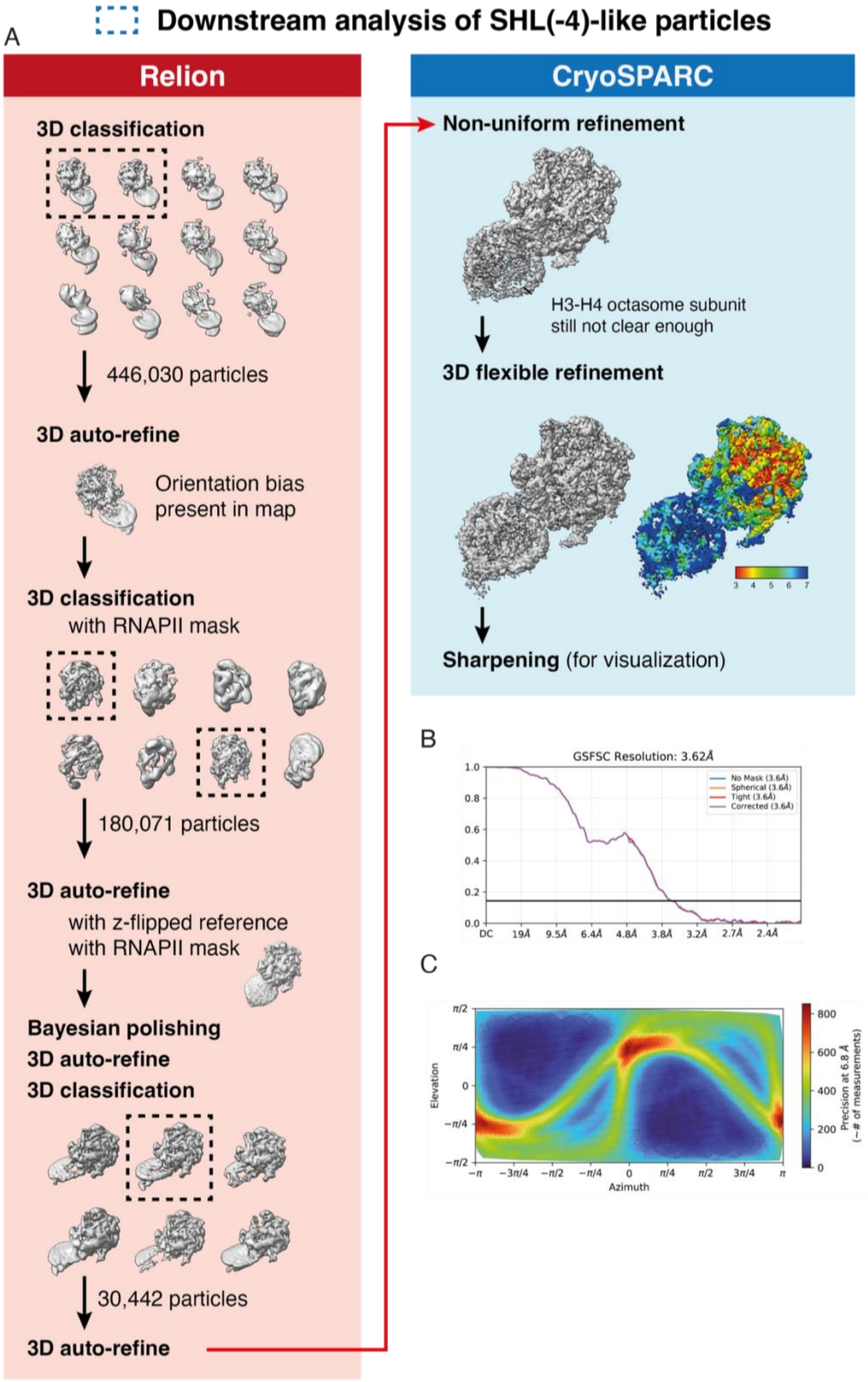

Downstream analysis of SHL(-4)-like particles
A
Relion
3D classification
446,030 particles
3D auto-refine
Orientation bias present in map
3D classification with RNAPII mask
180,071 particles
3D auto-refine with z-flipped reference with RNAPII mask
Bayesian polishing
3D auto-refine
3D classification
30,442 particles
3D auto-refine
CryoSPARC
Non-uniform refinement
H3-H4 octasome subunit still not clear enough
3D flexible refinement
Sharpening (for visualization)
B
GSFSC Resolution: 3.62Å
No Mask (3.6Å)
Spherical (3.6Å)
Tight (3.6Å)
Corrected (3.6Å)
DC 19Å 9.5Å 6.4Å 4.8Å 3.8Å 3.2Å 2.7Å 2.4Å
C
Elevation
Azimuth
Precision at 6.8 Å (~# of measurements)

**Supplementary Figure S5. Cryo-EM single particle analysis of the SHL(-4)-like particles.** (A) The SHL(-4)-like particles from Supplementary Figure 4 were further processed by both Relion and CryoSPARC. (B) Fourier shell correlation curves (C) Angular distributions

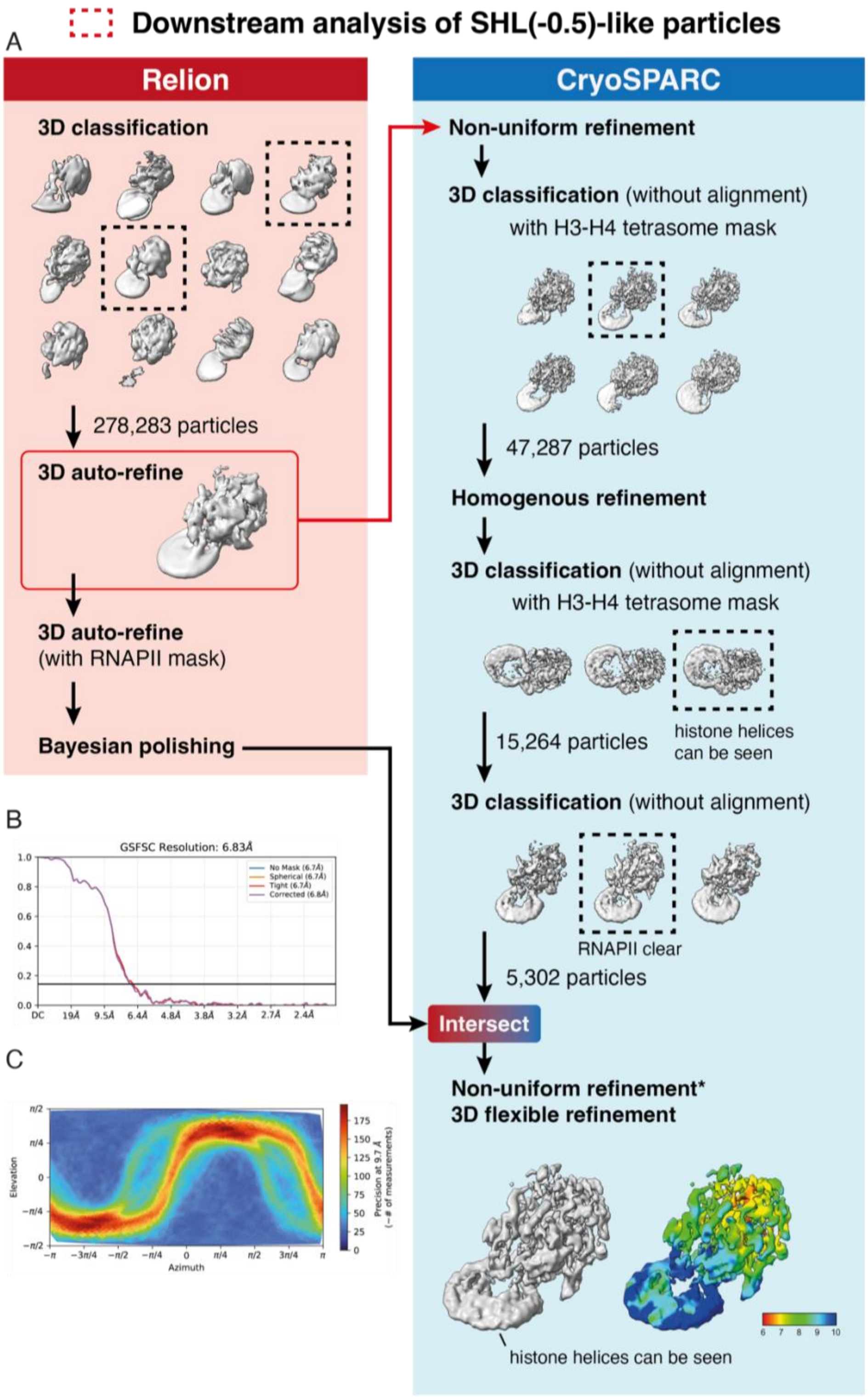


**Supplementary Figure S6. Cryo-EM single particle analysis of the SHL(-0.5)-like particles.** (A) The SHL(-0.5)-like particles from Supplementary Figure 4 were further processed by both Relion and CryoSPARC. (B) Fourier shell correlation curves (C) Angular distributions
